# Efficient Generation of *SOCS2* Knock-out Sheep by Electroporation of CRISPR-Cas9 Ribonucleoprotein Complex with Dual-sgRNAs

**DOI:** 10.1101/2024.06.30.601433

**Authors:** Ahmed K. Mahdi, Devon S. Fitzpatrick, Darren E. Hagen, Bret R. McNabb, Tara Urbano Beach, William M. Muir, Nicholas Werry, Alison L. Van Eenennaam, Juan F. Medrano, Pablo J. Ross

## Abstract

Knock-out (KO) sheep were produced using CRISPR-Cas9 ribonucleoprotein complexes in zygotes targeting an 85 bp section of the first exon of the Suppressor of Cytokine Signalling-2 (*SOCS2*) gene. Electroporation was performed 6 hours post-fertilization with dual-guide CRISPR-Cas9 ribonucleoproteins (RNPs). Fifty-two blastocysts were transferred to 13 estrus-synchronized recipients, yielding five live lambs and one stillborn. These lambs were all compound heterozygotes with mutations predicted to result in *SOCS2* KO. Three lambs carried large deletion alleles (259 bp, 1694 bp, and 2127 bp) that evaded initial detection via initial PCR screening. Off-target analysis identified a small number of mutations which may have been the result of off-target activity in regions with some homology to the guides, but notably such mutations were also observed in unedited controls. Further, we observed several orders of magnitude more mutations outside of these regions of homology in both edited animals and controls. Western blot and RT-PCR analysis of cell lines from SOCS2 KO lambs showed trace levels of *SOCS2* mRNA and SOCS2 protein. In conclusion, combining IVF and electroporation of dual-guide CRISPR-Cas9 RNPs was effective at generating KO sheep.

## 1. Introduction

Conventionally, genome editing reagents are introduced to livestock oocytes and embryos via microinjection; however, this approach requires expensive equipment, skilled personnel, and most importantly, frequently results in mosaicism ^1–4^, which can decrease the utility of producing genome-edited livestock by single-step approaches. Electroporation offers a rapid and efficient way to introduce editing reagents into mammalian zygotes. Electroporation is a commonly used method that can transfer different substances into a variety of cell types ^5–10^. Electroporation introduces short-term changes in plasma membrane permeability that allow for the transfer of CRISPR/Cas9 ribonucleoprotein (RNP) to the zygotic cytoplasm. Several reports have documented the production of gene-edited rats ^11^, mice ^12^, pigs ^13^, and cattle ^14^ by electroporation of zygotes in a single-step approach, as reviewed Lin and Van Eenennaam ^15^.

Suppressors of cytokine signaling (*SOCS*) are a family of eight proteins regulating cytokine intracellular signaling. One essential intracellular mediator of cytokine signaling is the Janus kinase and signal transducer and activator of transcription (JAK-STAT) pathway^16^. The *SOCS2* gene encodes a 198 amino acid (aa) protein with two functional domains: the central SH2 and the highly conserved C-terminal SOCS box. When growth hormone (GH) binds to its cell surface receptor, the JAK-STAT pathway transduces the signal throughout the cell and activates transcription of growth-related genes and *SOCS2*. SOCS2 proteins exert negative feedback on the growth hormone signaling pathway by binding to the growth hormone receptor (GHR) via the SH2 domain and blocking further phosphorylation and activation of STAT5 proteins. Additionally, SOCS2 functions as a substrate recognition subunit in the SOCS2 E3 ubiquitin ligase complex that tags the GHR for degradation^17, 18^. In *SOCS2* KO mice, an extended duration of STAT5 phosphorylation is observed in response to GH stimulation along with longer bones, 30-50% heavier mature body weight, and enlarged internal organs ^19–21^.

Many of the effects of *SOCS2* KO in mice are also observed in sheep (*Ovis aries*). In a study investigating genetic susceptibility to mastitis in dairy ewes, a naturally occurring nonsynonymous point mutation in *SOCS2* (p.R96C) was identified that disrupts the tyrosine binding pocket of the SH2 domain of SOCS2^22^. In vitro experiments showed a nearly complete lack of SOCS2 binding to its highest affinity phosphorylated residue on the GHR. This mutation was associated with increases of 24%, 18%, and 4.4% for height, weight, and milk yield, respectively, in *SOCS2* p.R96C homozygotes. However, there was also a strong positive correlation with lifetime somatic cell count score (a proxy for genetic susceptibility to mastitis)^22^. Recently, *SOCS2* p.R96C sheep were produced by programmable base editing^23^. In that study, 53 zygotes were microinjected and 4 lambs were born; one lamb was unedited, two had *SOCS2* indel mutations, and the third had primarily the intended single base pair substitution. However, bystander editing of nearby cytosine nucleotides occurred in all lambs and the total rate of intended substitution was below 45% for all animals, making the inheritance of the mutation inefficient using this method.

Previously, we successfully targeted *SOCS2* and *PDX1 genes* in sheep and *OTX2* in goat embryos by electroporating zygotes with long-high/short-low voltage pulses 6 hours after fertilization^24^. We found that this approach not only efficiently replaced microinjection, but also increased the biallelic mutation rate in small ruminant embryos. We further whole genome sequenced the genome-edited offspring and characterized the mutations present. In this study, we investigated the possibility of efficiently producing *SOCS2* knock-out (KO) sheep by embryo electroporation of dual gRNA Cas9/sgRNA RNPs designed to target an 85 bp region of exon 1 of *SOCS2* encompassing a part of the extended SH2 domain.

## 2. Materials and Methods

### 2.1. Experimental design

Two rounds of embryo production were used to generate *SOCS2* KO animals. In each round, zygotes were produced by in vitro fertilization of *in vitro* matured cumulus-oocyte complexes (COCs). Two sgRNA were introduced into the zygotes 6 hours after fertilization by electroporation with short-high/long-low voltage parameters as previously reported ^24^. Embryos were then cultured to the blastocyst stage and transferred to synchronized recipients. In total, we transferred 52 day 7 blastocysts to 13 estrus synchronized recipients, in a ratio of 4 blastocysts per recipient. Pregnancies were checked by ultrasound 30 days after fertilization. Lambs were genotyped with a combination of PCR-Sanger sequencing and whole genome sequencing. To test *SOCS2* functionality, RT-PCR was used to detect the transcription of *SOCS2* sequences and western blotting (WB) was used to detect *SOCS2* protein production in live animals.

### 2.2. sgRNAs design

Two sgRNAs were designed using the online E-CRISP gRNA design tool (http://www.e-crisp.org) and ordered from Synthego (Synthego Corporation, Redwood City, CA, USA) to target the beginning and end of the sheep *SOCS2* exon 1 (Figure 1 and Supplementary Table 1); the expected deletion was 85 bp long. We used the Oar_v3.1 reference genome (GCA_000298735.1) ^25^, and strict search criteria specifically for Knockout.

**Figure 1.**
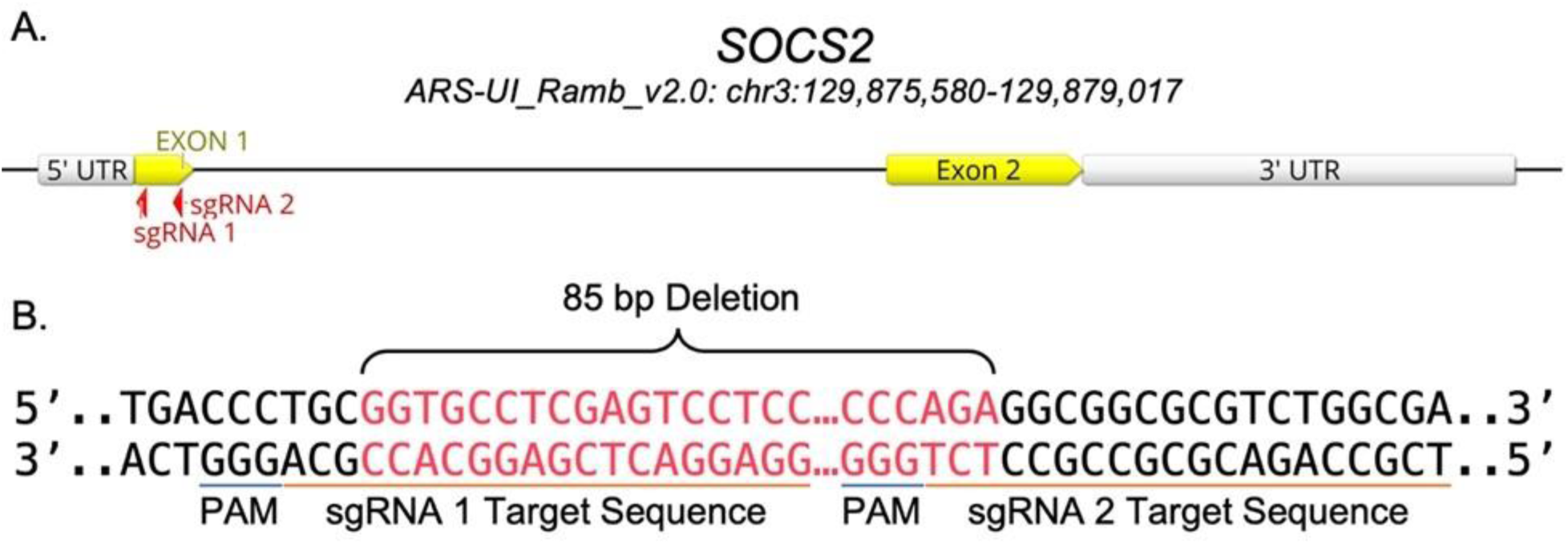
(A) *SOCS2* gene and sgRNA target loci. (B) sgRNA 1 and 2 target sequences and predicted deletion between target loci.

### 2.3. Gametes

#### 2.3.1. Cumulus Oocytes Complexes (COCs)

sheep ovaries were collected from a local abattoir (Superior Farms, CA) and transported to the laboratory in 37° C saline. A 21 G butterfly needle aspirated COCs after applying vacuum pressure.

### 2.4. In vitro Embryo Production

#### 2.4.1. In vitro oocyte maturation

good quality COCs with several cumulus cell layers, and homogenous cytoplasm were selected for maturation. COCs were washed five times with warm washing media and then transferred in groups of 50 to 400 µL of maturation medium (BO-IVM, IVF BIOSCIENCE) and incubated for 22-24 h at 38.5 °C, 5% CO2, 5% O2, 90% N2 and saturated humidity.

#### 2.4.2. In vitro fertilization

Matured COCs were washed three times with warm and pre-equilibrated fertilization media (BO-IVF, IVF BIOSCIENCE) and transferred in groups of 50 to 400 µl of fertilization media. Semen was washed twice by centrifugation at 328 g for 5 minutes with 2 mL of semen preparation media (BO-Semen Prep, IVF BIOSCIENCE). The sperm concentration was adjusted with fertilization media to 2 x 10^6^ spermatozoa/ml and 50 µl added to COCs for 6 hours 38.5 °C, 5% CO2, 5% O2, 90% N2 and saturated humidity.

#### 2.4.3. In vitro culture

Cumulus cells were stripped off presumptive zygotes by vortexing for 3 minutes in SOF-HEPES media, then zygotes were washed five times, and transferred in groups of 50 to 500 µL of culture media (BO-IVC, IVF BIOSCIENCE) covered with 400 µl mineral oil and incubated for seven days at 38.5° C, 5% CO2, 5% O2, 90% N2, and saturated humidity.

### 2.5. Ribonucleoprotein (RNP) Preparation

All RNA work was done in an RNAse-free workstation at room temperature. 40 ng/uL of each sgRNA was mixed with Cas9 protein (PNA Bio) in mass ratio of 1:2 and incubated on ice for 10 minutes, RNPs were mixed immediately prior to electroporation.

### 2.6. Electroporation

Electroporation was performed using the NEPA21 Super Electroporator in 1 mm Nepa Electroporation Cuvettes (EC-001). All work was done in an RNAse-free workstation. Denuded zygotes were washed three times with SOF-HEPES and three times with warm Opti-MEM (Thermo Fisher Scientific #31985062). Then zygotes were moved to a new drop of Opti-MEM and mixed with RNPs to a total volume of 20 µL, loaded into an electroporation cuvette. The poring parameters were set to four unipolar electric pulses of 40 volts of 3.5 msec each and the transfer parameters were set to five bipolar electric pulses of 5 volts for 50 msec each. After delivery of electric pulses, the cuvette was washed several times with SOF-HEPES, and the recovered zygotes were washed three times with SOF-HEPES and one time with BO-IVC before being moved to pre-warmed and pre-equilibrated BO-IVC media and cultured for seven days.

### 2.7. Sheep Embryo Transfer

#### 2.7.1. Recipient Selection and Estrus Synchronization

All experiments involving animals were approved and performed in accordance with the University of California Davis Institutional Animal Care and Use Committee (IACUC Protocol # Protocol #: 20259). Healthy ewes were estrus synchronized using an intravaginal progesterone device (0.3 g of progesterone; CIDR-G; Zoetis). CIDRs were inserted for six days. On the day of CIDR removal, prostaglandin F2-alpha analog (10 mg dinoprost thrometamine; Zoetis) and PG600 (400 IU PMSG, 200 IU hCG; Intervet) were injected intramuscularly. Estrus was detected every 12 hours after removing the CIDRs with the assistance of a vasectomized ram for heat marking. Thirteen ewes were selected for the embryo transfer 6 days post-estrus.

#### 2.7.2. Laparoscopic Embryo Transfer

Food and water were removed 24 and 12 h prior to surgery, respectively. Fifteen minutes before surgery, recipients were sedated by administration of 1.1-2.2 mg/kg of ketamine and 0.2-0.3 mg/kg of Midazolam. After preparing the surgery site, 2% Lidocaine was injected at the incision site to induce local anesthesia. The corpus luteum (CL) was visualized by laparoscopy (Karl Storz, Germany), and four blastocysts were injected into the tip of the uterine horn ipsilateral to the CL.

#### 2.7.3. Pregnancy Diagnosis

Pregnancy diagnosis was performed 30 days post-estrus by abdominal ultrasonography. The frequency of the transducer was 3.5 MHz.

### 2.8 Genotyping

#### 2.8.1 Genomic DNA extraction

Tissue/blood was collected from genome edited lambs and two control rams, and DNA was extracted using DNeasy Blood & Tissue Kit (Qiagen Cat. No. / ID: 69504).

#### 2.8.2 In vitro On-Target Indel Detection

PCR reactions were prepared with 10 μL of Promega GoTaq Green Master Mix (2x), 1 μL forward primer (10μM), 1 μL reverse primer (10μM), 100 ng of genomic DNA, and molecular grade nuclease-free water to a final volume of 20 μL. CRISPR-Cas9 target region was amplified by PCR, primer sequences and amplification conditions are available in Supplementary Table 2. PCR products were run on 1% or 2% agarose gels prepared with 60 or 120 mL of 1% tris base acetic acid and EDTA (1% TBE), and SYBR^TM^ Safe gel stain (10,000X) diluted to 6X. Visualization of bands was performed under blue and UV light and compared to a control amplicon. DNA fragments were cut out of gels individually using a scalpel and purified PCR products were gel-extracted by either by the QIAquick Gel extraction kit (QIAGEN) or a freeze and squeeze method of placing the gel on top of a filter tip in a 1.5 mL conical tube, freezing the sample for 5’ at -80 °C then centrifuging for 3 minutes at 14,000 RPM. Gel-purified PCR products were outsourced to GeneWiz (Azenta, South Plainfield, New Jersey) for purified PCR product Sanger sequencing. To detect indels, first, each amplicon was aligned to a control amplicon using SnapGene software (version 6.1.1), to visualize the modifications. The sequences were checked for insertions, deletions, and double peaks.

### 2.09. Allele Analysis

The consequence of each mutant allele was assessed with ENSEMBL Variant effect predictor (release 108). For prediction of the effects when two variants were associated within the same allele present in target site 1 for female, tag #7181, and male tag #7186 all lambs were assessed for the effect of the first mutation, due to a current limitation of the software to only analyze the effect of a single variant. Missense mutations that also resulted in frameshifts were not identified as high impact frameshifts, but as coding variants.

### 2.10. Whole Genome Sequencing

Genomic DNA from the six genome edited lambs that were born, and their two unedited sires (Ram 1 and Ram 2) were used for whole genome sequencing (WGS) with the Illumina NovaSeq platform following manufacturer instructions. Insert sizes were approximately 300 bp with 2x150 bp paired-end reads to an approximate depth of 15-46x per sample. Qualified reads were mapped to the ARS-UI_Ramb_V2.0 (GCA_016772045.1) ^26^ and Oar_v4.0 (GCA_000298735.2) reference genomes using Burrows-Wheeler Aligner-Maximum Exact Match (BWA-MEM) and indexed with BWA-Index (BWA tools v7.1). Aligned WGS BAM files were visualized with Integrative Genome Viewer (v2.13.2).

### 2.11. Single Nucleotide Variants (SNV) and small indel identification

For a detailed analysis, and genome wide identification of SNVs, the Illumina WGS data were mapped to the reference genome (ARS-UI_Ramb_V2.0) using CLC Genomic Workbench V11.0 (https://digitalinsights.qiagen.com). the reads were mapped to the reference by using conservative mapping parameters which required that 75% of a read to map uniquely to the reference. Further affine gap cost penalty was used for insertions and deletions to the maximum cost allowed of 6 to open an insertion or deletion position and an extension cost of 1. Following mapping, all variants, including SNVs, small insertions, and deletions (INDELs < 30bp) were called using the basic variant detection tool using the following parameters: only reads mapped as intact pairs were included, with a minimum Pfred quality score of 30 for the central variant, as well as the 5 adjoining bases on either side. The final step was to require a minimum variant frequency of 25% within a sample.

### 2.12. Structural variant (SV) analysis

Genome wide SV (deletions, inversions, intra-chromosomal translocations, tandem duplications), were identified by mapping reads to the reference with more liberal parameters than for SNV detection in order to minimize rejection of reads due to misalignment caused by large INDELS. CLC genomics software checks read mappings for evidence of breakpoints using “unaligned end” signatures. “Unaligned end” refers to the end of a read that do not map to the reference sequence at the positions presented in the read after the portion that does align to the reference. Thus, to maximize discovery of SV, and to minimize rejection of reads due reference misalignment, a minimum of only 10% of a read was required to be mapped uniquely to the reference, allowing any unaligned ends to be incorporated in a stringent search for SV. The algorithm first identifies positions in the mapping(s) with an excess of reads with left (or right) unaligned ends. Once these positions and the consensus sequences of the unaligned ends are determined, the algorithm maps the determined consensus sequences to the reference sequence around other positions with unaligned ends. If mappings are found that are in accordance with a ’signature’ of a structural variant, a structural variant is called.

### 2.13 On Target analysis

In addition to the SV analysis using CLC, a manual confirmation method was also used as follows: For deletions that appeared to be within the span of overlapping paired reads the following method was used: A) based on visual examination of the unaligned ends, haplotypes were identified and defined by a portion of the sequence in the unaligned ends plus adjoining sequences that aligned with the reference. These haplotypes were extracted from the raw fastq data using custom scripts and stored in separate files. B) the haplotype-resolved reads were then aligned using the DeNovo aligner of CLC genomics. The resulting consensus alignment of each haplotype was compared to the *Ovis aries rambouillet* reference genome (ARS-UI_Ramb_V2.0) using NCBI’s basic local alignment search tool (BLAST). Alignments that were not contiguous with the reference were flagged as possible SVs. C) The actual SV was identified by matching the contiguous parts of the BLAST with the reference sequence and counting the number of base pair (bp) deletions or insertions needed to enable alignments of the fragments to reference sequence.

For deletions that were greater than the span of overlapping paired reads, an alternative manual method of verification was employed. This method utilized paid reads, which are fragments with ends that were sequenced as a physical unit, but were mapped to the reference as broken reads, i.e. each end mapped to a different region. By visual examination, reads that were broken but at least partially mapped to the reference in the target region, were extracted and their read IDs tabulated. The mates to the extracted reads were found by searching the raw fastq data file for each mate’s matching ID. The extracted reads from the target region along with their mates were then aligned using the DeNovo aligner and the process proceeded as previously described for short deletions.

### 2.14. Off-Target analysis

Potential off-target binding sites with up to 4 bp mismatches (Supplementary Table 3) were examined by mapping single nucleotide variants (SNV) and large structural variants (SV) identified by CLC spanning ±100 bp away of the putative cut site. To investigate whether the identified variants were associated with potential off-target activity, we developed a custom R script which detects if the possible binding site is within the mutated region or within an extended range spanning an additional 100 bp from either end (R version 4.3.2). Chi-square tests were conducted using Monte Carlo simulated p.values, compare the frequency of mutated regions which contained a putative off-target site between edit samples and wild-type control rams.

### 2.12. Fibroblast cultures

Fibroblast cell lines from lambs were derived as previously described ^27^. Briefly, 4-mm skin biopsies were taken from two different sites in each lamb. The skin was dissected, and small dermal pieces were cultured for ten days in DMEM high glucose media (Thermo Fisher, Cat#11995065) supplemented with 20% FBS (Gibco Cat# 10100139), 1% GlutaMAX (Gibco, Cat#35050061), and 1% Penicillin-Streptomycin (Gibco, Cat#15140122). Cells were passaged using trypsin at least two times to obtain homogenous fibroblast cell lines.

### 2.13. Two-step Real-Time RT-PCR

RNA was extracted from fibroblasts following the RNeasy Mini Kit (Qiagen Cat#74104) protocol. Extracted RNA was quantified using the NanoDrop 2000C Spectrophotometer (ThermoScientific), and its integrity was assessed by 1.5% agarose gel electrophoresis. 50 ng of the extracted RNA was reverse transcribed using a High-Capacity cDNA Reverse Transcription Kit (Thermo Fisher Cat#4368813). RT-PCR was performed via QuantStudio 3 (Applied Biosystems) in 20 µL reactions containing 10 µL of Power Track™ SYBR Green Master Mix (Thermo Fisher Cat#A46012), 0.2 µL of forward and 0.2 µL of reverse primer (10 µM stock of each primer, Supplementary Table 2), 2 µL of the cDNA sample (10 ng/µL), and 7.6 µL of water. Each sample was run in duplicate for each independent cell line.

### 2.14. Protein Extraction and Western Blotting

Proteins were extracted from fibroblast cells using RIPA Buffer (Sigma #R0278) with protease inhibitors (Roche #11836153001). Proteins were quantified using Pierce™ BCA assay (Thermo Fisher Scientific, Cat#23225). Western blotting was performed as previously described^28^. 20 µg of proteins per sample were electrophoresed on 12% Mini-PROTEAN TGX precast protein gels (Bio-Rad) for 5 minutes at 50 volts followed by 115 minutes at 100 volts. Proteins were transferred to Immun-Blot PVDF membrane (Bio-Rad) for 120 minutes at 100 volts then blocked for one hour in 3% BSA TBS-T buffer. The membrane was incubated at 4°C overnight with the following primary antibodies anti-SOCS2 antibody (Abcam#ab66733,1:1000) and anti-β-Actin antibody (C4) (Santa Cruz#sc-47778,1:1000). Then the membrane was washed three times in TBS-T and incubated at room temperature for 2 hours with the following secondary antibodies goat anti-rabbit IgG H&L (HRP) (Abcam #ab205718, 1:2000) and m-IgG Fc BP-HRP (Santa Cruz, #sc-525409,1:2000). The ChemiDoc-It imaging system was used to visualize the membrane.

### 2.15. Preliminary Phenotype of Homozygous SOCS2 F_0_ KO lambs

The growth variables of weight, height, and length were measured in the three SOCS2(-/-) KO founder sheep (7181 female, 7182 & 7183 male) through 20 weeks and compared to two phenotype control ewes in the flock that were born at a similar time and raised in the same pen under commercial conditions.

## 3. Results

### 3.1. Pregnancies, Abortions, Lambing, and Survival

Of the thirteen recipients that received embryos, six were determined by ultrasound to be pregnant at day 30. Two recipients aborted at weeks 13-14 of pregnancy. One recipient delivered a stillborn male lamb (#7186), which may have died because of its large size (12.2 kg). Another recipient gave birth to triplets, two males and one female. The female (#7184) was euthanized at 2 days of age due to breathing difficulties, one male (#7185) died at 5 days of age due to while the other male was healthy (#7183). Two recipients gave birth to one healthy lamb each, one male (#7182) and one female (#7181) (Figure 2).

**Figure 2.**
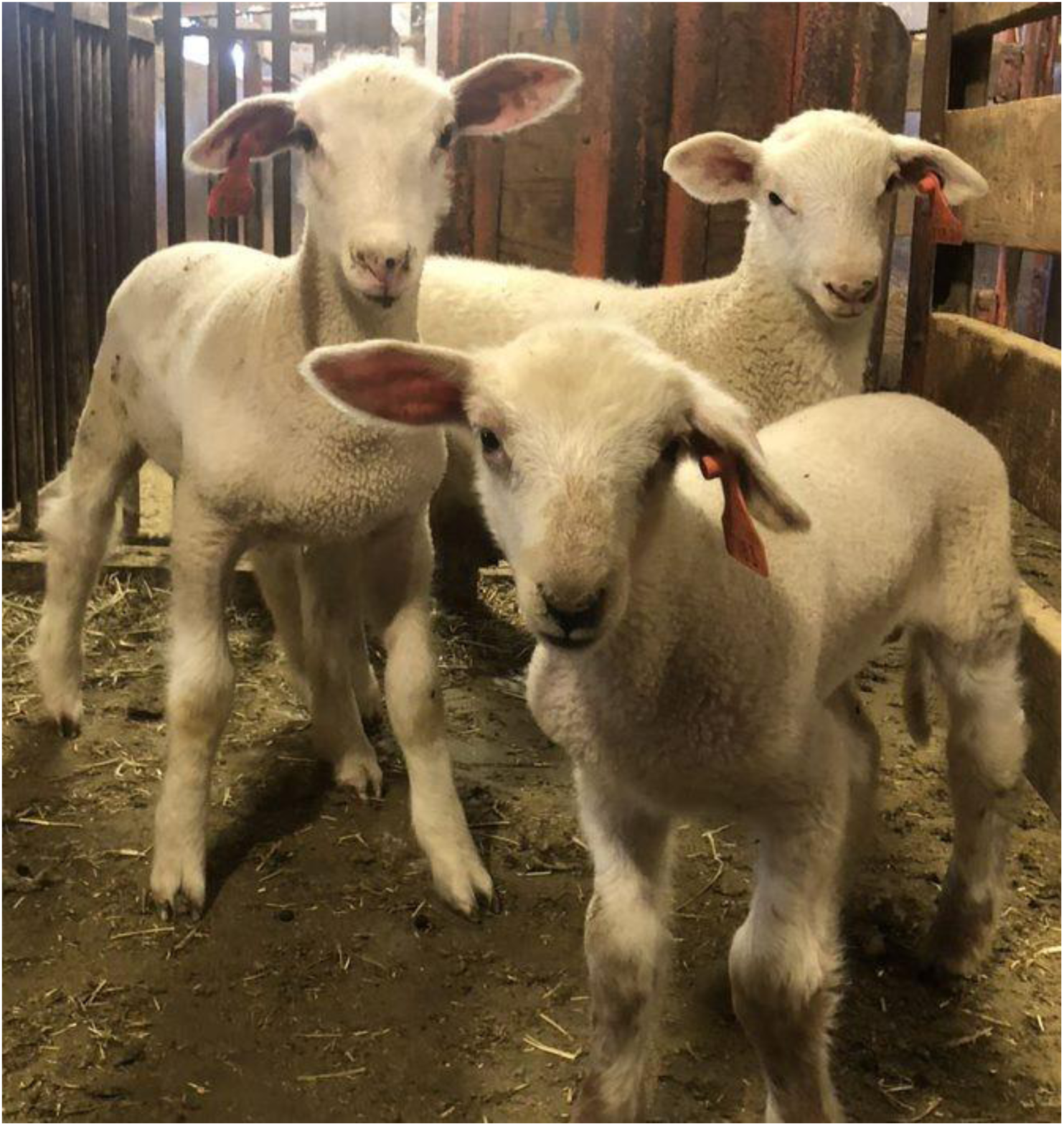
SOCS2 genome edited lambs female tag #7181 (front), male tag #7182 (left), and male tag #7183 (right).

### 3.2. Genotyping Analysis of Genome Edited Lambs

Genotyping of the genome edited offspring was initially performed by PCR amplification of a 479 bp region flanking the two target sites. All genome edited lambs appeared to have deletion alleles via gel electrophoresis (Supplementary Figure 1). Analysis of WGS data revealed that both sgRNA 1 and sgRNA 2 target sites were disrupted in each of the 6 genome edited lambs (Figure 3). Deletions varied in size from 30 bp in an allele of female tag #7184 to 2127 bp in a female tag #7181 allele. Three large deletions in the healthy lambs: female tag #7181, male tag #7182, and male tag #7183, evaded initial detection by PCR followed by Sanger sequencing due to the elimination of one or multiple primer binding sites. Additionally, three intermediate-sized deletions of 30 bp, 101 bp, and 109 bp were identified, with the 101 bp deletion present in male tag #7188 spanning both target sites and the other two initiating at target site 2. None of the mutations were the exact 85 bp deletion between the predicted cleavage sites, but 5 deletions were within a 5 bp window of both cut sites (Figure 3). Finally, four small insertions were observed, with a single base pair guanine insertion at cut site 1 occurring in allele 1 of the female tag #7181 and allele 1 of stillborn male tag #7186, as well as a “G” insertion and “TTT” insertion occurring in tag #7186 allele 2 at cut sites 1 and 2, respectively (Figure 4).

**Figure 3.**
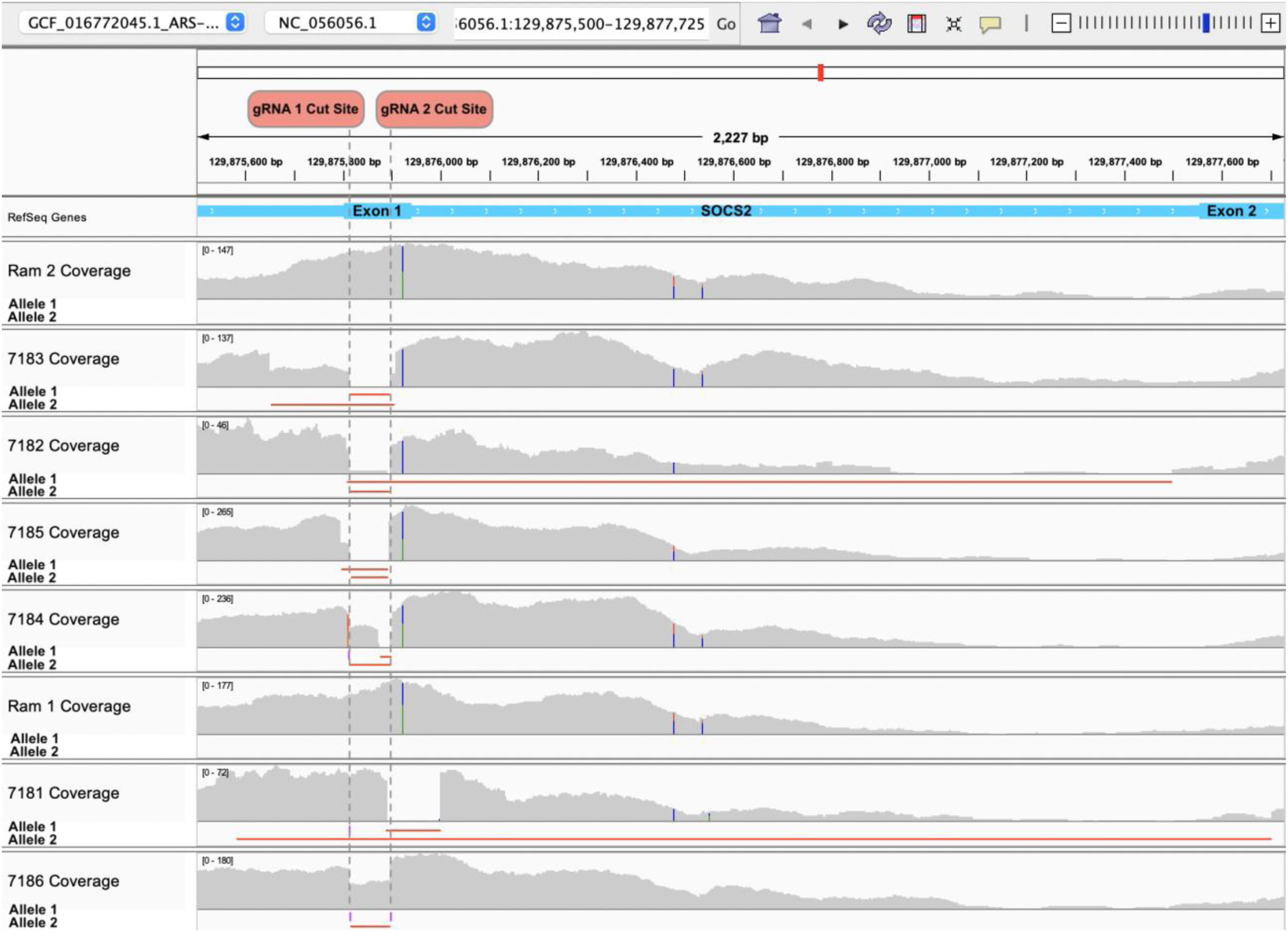
Read coverage tracks at the SOCS2 target locus for Ram 2, Ram 2’s genome edited offspring tag #7183, #7182, #7185, and #7184; Ram 1, and Ram 1’s genome edited offspring tag #7181 and #7186. The read depth is indicated in gray on the read coverage track, the span of each deletion is indicated in red on the allele tracks, and insertions are indicated as purple rectangles on the allele tracks.

**Figure 4.**
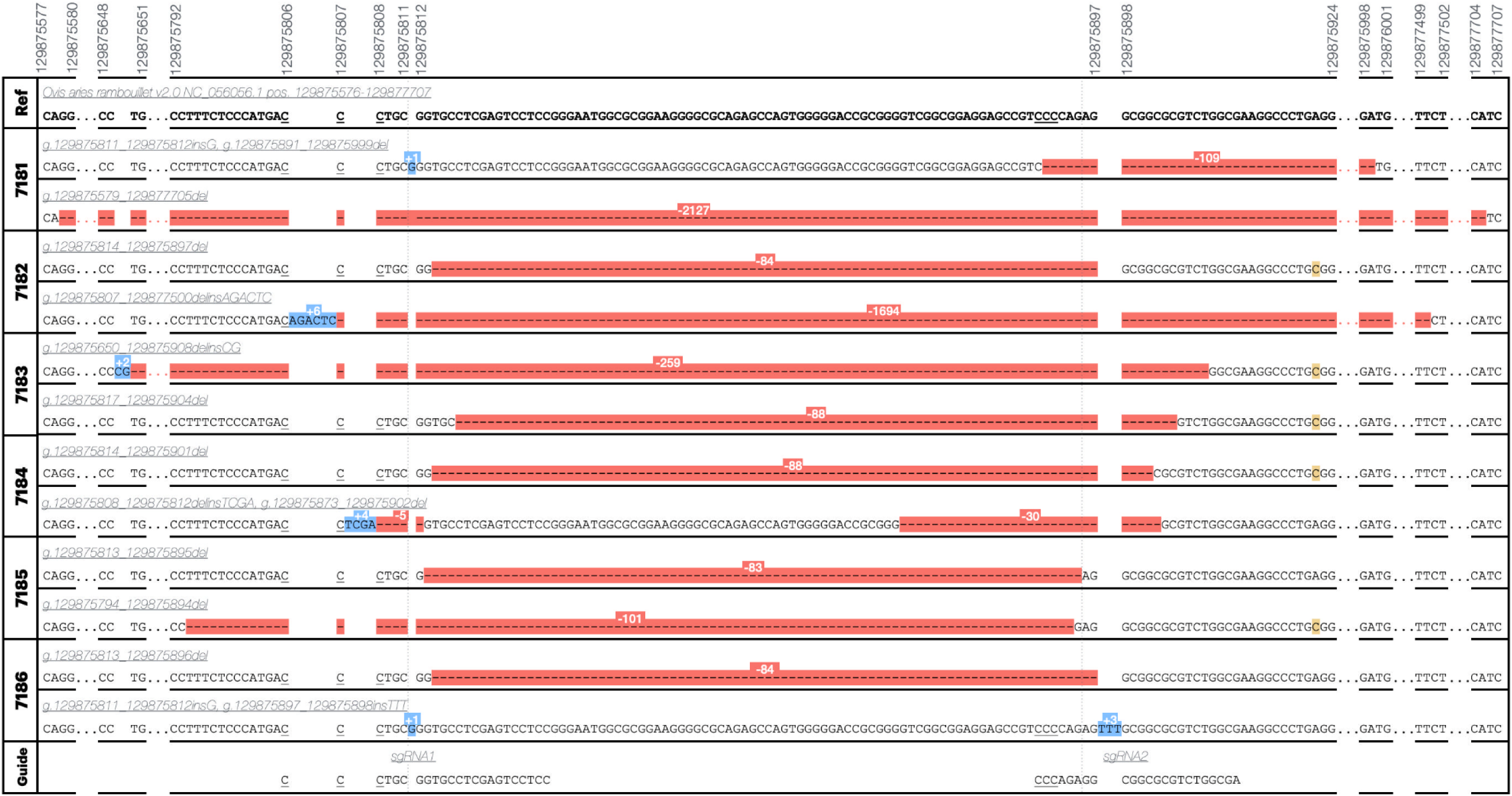
CRISPR/Cas9 genome edited alleles. Variant names are according to human genome variation society nomenclature guidelines for genomic variants and indicate positions on chromosome 3 (NC_056056.1) in the ARS-UI_Ramb v2.0 reference genome (ref). Red-highlighting indicates a deletion, blue an insertion, yellow an endogenous SNP not considered to be a result of genome editing. Ranges not shown are indicated by ellipses and are either wild-type sequence or continued deletion sequence. PAM sites (NGG reverse-complement) are indicated by underlining.

A small number of mosaic reads were observed in the WGS data, but no wild type reads spanning both cut sites were observed and all mosaic reads were present at <5% minor allele frequency (MAF) and may be artifacts of mismapped reads or index hopping, which according to the manufacturer occur at low rates in samples.

The predicted effect of each of the major mutations in the was analyzed by prediction of the resulting *SOCS2* translation (Table 1). All analyzed variants were predicted to result in knock out of the *SOCS2* gene either through deletion of a large section of the protein, deletion of the start site, or a frameshift and premature introduction of a stop codon.

**Table 1.** Predicted amino acid sequences per edited allele. Variant names indicate variant position on chromosome 3 (NC_056056.1) in the ARS-UI_Ramb v2.0 reference genome. Predicted amino acid sequences were generated by *in silico* translation on SnapGene: bold text indicates amino acid sequence from endogenous start site, asterisk indicates stop codon.

| Sample | Variant Name | Predicted Amino Acid Sequence | Mutation |
| --- | --- | --- | --- |
| Wild-type | *SOCS2* | **MTLR**CLESSGNGAEGAQSQWGTAGSAEEPSPEAARLAKALRELSHTGWYWGNMTVNEAKEKLKEAPEGTFLIRDSSHSYLLTISVKTSAGPTNLRIEYQDGKFRLDSEICVKSCLKQFDSVVHLIDYYVQMCKDKRTGPEAPRNGTVHLYLTKPLYTSAPPLQHLCLRTINKCTSTIWGLPLPTRLKDYLEEYKFQV* | N/A |
| 7181 | *g.129875811_129875812insG, g.129875891_129875999del* | **MTLR**VPRVLREWGRGAEPVGDGRGVGGGAVVGTGEI* | Frameshift, nonsense |
| 7181 | *g.129875579_129877705del* | MCQVQA* | Original start deleted, frameshift, nonsense |
| 7182 | *g.129875814_129875897del* | **MTLR**AARLAKALRELSHTGWYWGNGMTVNEAKEKLKEAPEGTFLIRDSSHSYLLTISVKTSAGPTNLRIEYQDGKFRLDSEICVKSCLKQFDSVVHLIDYYVQMCKDKRTGPEAPRNGTVHLYLTKPLYTSAPPLQHLCLRTINKCTSTIWGLPLPTRLKDYLEEYKFQV* | 28 AA deletion |
| 7182 | *g.129875807_129877500delinsAGACTC* | **MTLR**CESVWRRP* | Frameshift, nonsense |
| 7183 | *g.129875650_129875908delinsCG,* | MTVNEAKEKLKEAPEGTFLIRDSSHSYLLTISVKTSAGPTNLRIEYQDGKFRLDSEICVKSCLKQFDSVVHLIDYYVQMCKDKRTGPEAPRNGTVHLYLTKPLYTSAPPLQHLCLRTINKCTSTIWGLPLPTRLKDYLEEYKFQV* | Original start deleted, 52 AA deletion |
| 7183 | *g.129875817_129875904del* | **MTLR**CVWRRP* | Frameshift, nonsense |
| 7184 | *g.129875814_129875901del* | **MTLR**RVWRRP* | Frameshift, nonsense |
| 7184 | *g.129875808_129875812delinsTCGA, g.129875873_129875902del* | **MT**SSASSPPGMARKGRRARGDRGASGEGPEGTQSHRLVLGKYDC* | Frameshift, nonsense |
| 7185 | *g.129875813_129875895del* | **MTLR**GGASGEGPEGTQSHRLVLGKYDC* | Frameshift, nonsense |
| 7185 | *g.129875794_129875894del* | MTVNEAKEKLKEAPEGTFLIRDSSHSYLLTISVKTSAGPTNLRIEYQDGKFRLDSEICVKSCLKQFDSVVHLIDYYVQMCKDKRTGPEAPRNGTVHLYLTKPLYTSAPPLQHLCLRTINKCTSTIWGLPLPTRLKDYLEEYKFQV* | Original start deleted, 52 AA deletion |
| 7186 | *g.129875813_129875896del* | **MTLR**AARLAKALRELSHTGWYWGNGMTVNEAKEKLKEAPEGTFLIRDSSHSYLLTISVKTSAGPTNLRIEYQDGKFRLDSEICVKSCLKQFDSVVHLIDYYVQMCKDKRTGPEAPRNGTVHLYLTKPLYTSAPPLQHLCLRTINKCTSTIWGLPLPTRLKDYLEEYKFQV* | 28 AA deletion |
| 7186 | *g.129875811_129875812insG, g.129875897_129875898insTTT* | **MTLR**VPRVLREWGRGAEPVGDGRGVGGGAVPRVCGASGEGPEGTQSHRLVLGKYDC* | Frameshift, nonsense |

### 3.3 Off-Target Analysis

Genomic regions with up to 4 bp mismatches to the target sequence were determined to identify 133 possible off-target sites of the two guide RNAs (Supplementary Table 3). Cross-referencing these possible off-target sites to genomic regions where a mutation occurred (± 100bp) as compared to the reference genome (Ovis aries rambouillet.ARS-UI Ramb v2.0) revealed a total of 40 mutations across all samples. Of these, 6 of these putative off-target sites were directly within the mutated region itself (Supplementary Table 4). Chi-square analysis indicated that off-target sites were no more likely to occur within 100 bp of a mutation in edited sheep as compared to wild-type controls (p = 0.7076) (Figure 5). Additionally, mutations occurred outside of this region at a rate that is several orders of magnitude greater in both edited animals and controls.

**Figure 5.**
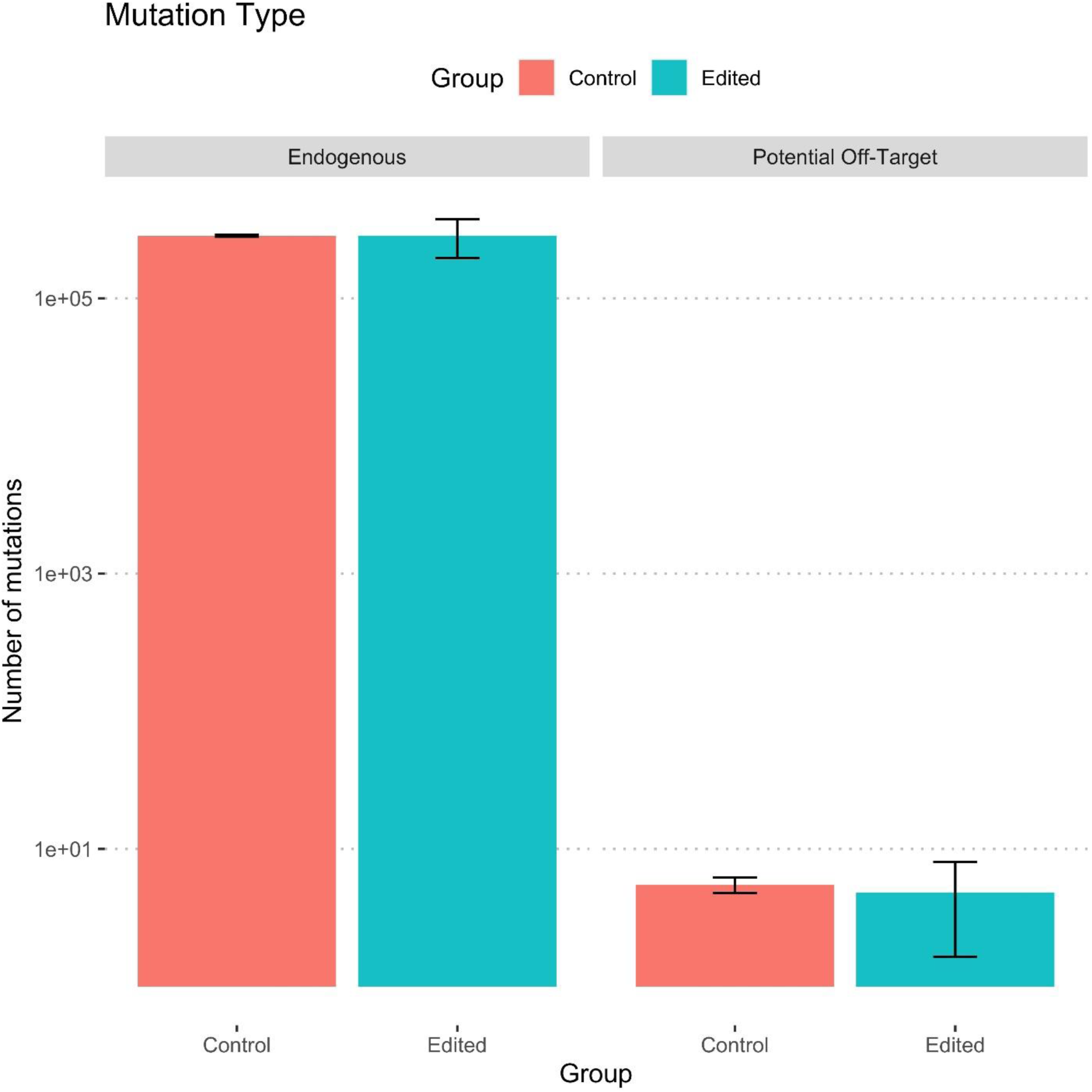
Boxplot comparison of mutated regions to assess potential off-target activity. Endogenous mutations occurred with no potential off-target binding sites (with up to 4 mismatches) occurring within 100 bp from the mutated regions. Potential Off-target indicates there was a mismatched binding site within 100bp of the mutated region. A Chi-square test did not show a significant difference between the frequency of potential off-target edits occurring in control animals (Ram 1, Ram 2; n=2) as compared to edited animals (7181-7186; n=6), p = 0.7076.

### 3.4 Gene Expression Results

The *SOCS2* mRNA levels were decreased in males tag #7182 and male tag #7183, and female tag #7181, respectively, compared to wild-type (Supplementary Table 5). Trace levels of SOCS2 protein expression were detected in a western blot (Figure 6).

**Figure 6.**
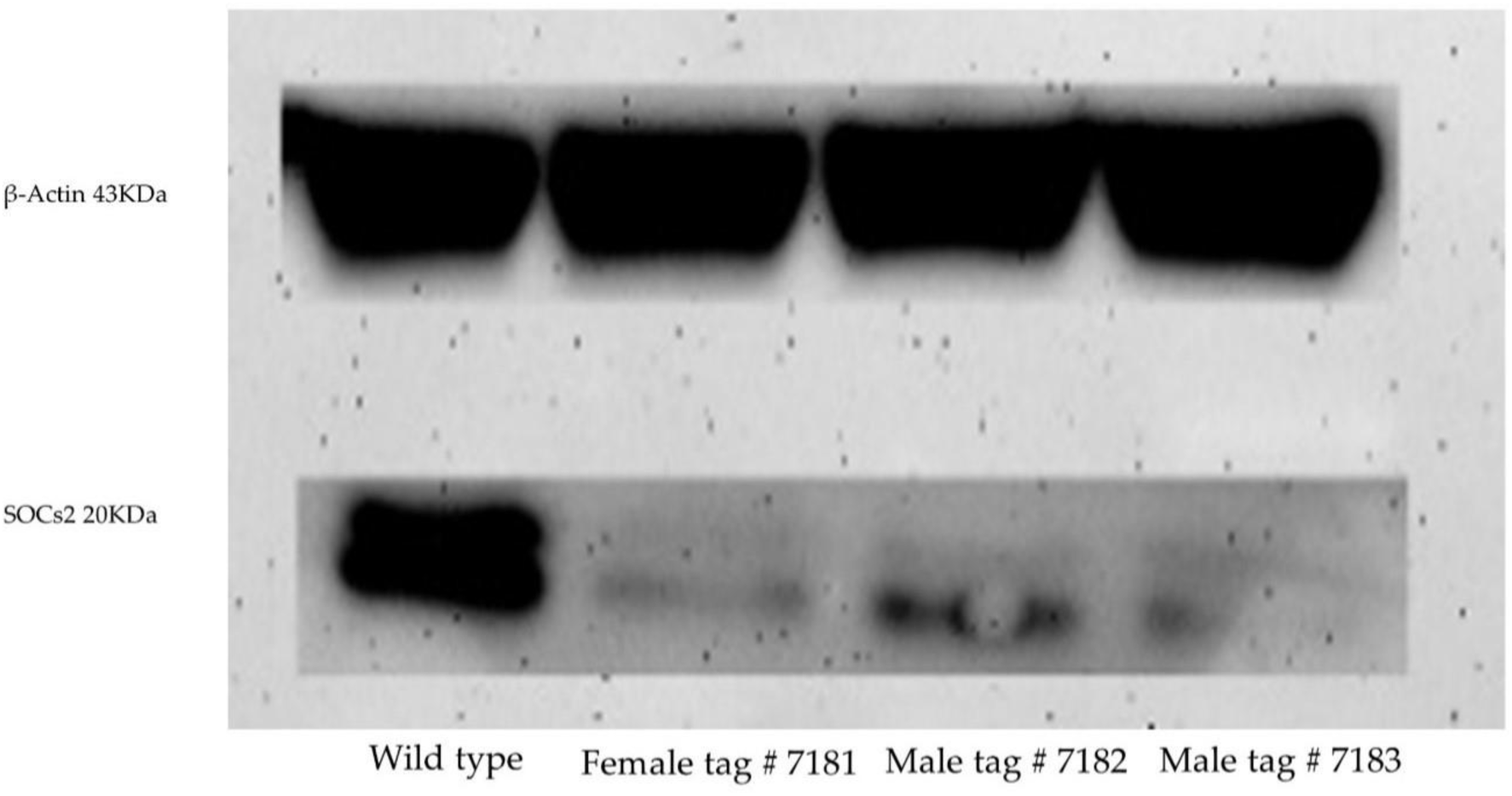
Western blot of protein extracted from fibroblast cell lines derive from three healthy *SOCS2* knock-out lambs produced by zygote electroporation of CRISPR-Cas9.

### 3.5 Preliminary Phenotype of Homozygous *SOCS2* F_0_ KO lambs

Preliminary results showed that the *SOCS2*(-/-) KO sheep had an average weight of 66.4 kg at 20 weeks of age compared to 42.3 kg for the controls. It is important to note that both phenotype control sheep were female, and 7183 was the only surviving lamb from a set of triplets. Furthermore, the *SOCS2*(-/-) KO sheep reached the final weight of the control sheep in approximately half the time. The increase in weight was accompanied by an approximately proportional increase in height and length (Figure 7).

**Figure 7.**
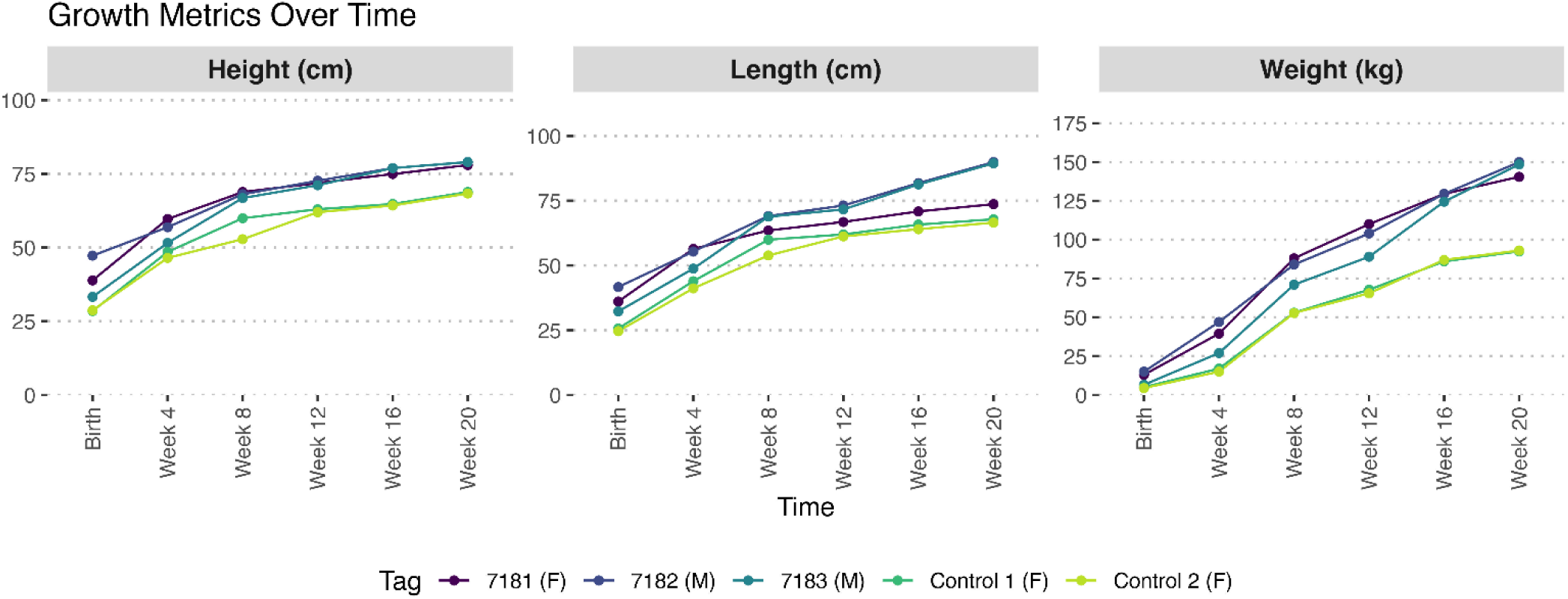
Preliminary growth results of the founder F_0_ *SOCS2* genome edited knock-out lambs (tag #7181 female, 7182 male, 7183 male) and two phenotype control ewes over the first 20 weeks.

## 4. Discussion

Electroporation of Cas9 RNP is a recent advance in the genome engineering of livestock embryos that allows for batched introduction of genome editing reagents for up to around 100 oocytes or embryos simultaneously, which offers several advantages over individual oocyte microinjection^15^. Early evidence suggests that electroporation is highly efficient at creating genome edited embryos with a minor effect on embryo survival so long as electroporation is performed at the time of fertilization or at the pronuclear stage and electroporation settings are optimized for the species of interest ^15, 24^. In mice, electroporation of gRNA into Cas9 expressing embryos was shown to result in a 100% or close to 100% mutation efficiency with no mosaicism^29^. Previous studies, using microinjection of CRISPR-Cas9 into sheep zygotes reported a lower mutation efficiency (10-38%) ^30–32^.

Our previous work ^24^ showed that electroporation was superior to microinjection in targeting the sheep *SOCS2* gene in embryos. Electroporation generates about 2 million of resealable pores in the plasma membranes ^33^ which compares to a single-entry point for microinjection approaches. The high efficiency of electroporation reduces the number of zygotes required to produce genome edited animals^13^, increasing the efficiency of any potential genome modified large animal production system. Electroporation does not require expensive equipment and high technical proficiency facilitating the wide spread use of gene editing technology^34^. Furthermore, electroporation has the advantage of simultaneous delivery of CRISPR-Cas9 RNPs to many embryos, which is not possible with microinjection where each embryo needs to be processed independently. In our hands, electroporation allowed processing up to 500 zygotes per hour in comparison to 40-50 per hour when microinjecting.

In the current study, we report the generation of *SOCS2* knock-out sheep by CRISPR-Cas9 electroporation of dual sgRNAs into *in vitro* produced zygotes. We obtained six lambs, of which three survived to 20 weeks. We electroporated the zygotes 6 hours after *in vitro* fertilization to increase the chances of biallelic non-mosaic mutations by delivering the CRISPR-Cas9 reagents to the embryo before the DNA replication preceding the first embryonic cleavage (4). The method and the timing of editing reagent delivery are important, as electroporation of mouse zygotes before E0.5 has been shown to decrease mosaicism ^1^ and electroporation of sheep zygotes 8 hours after fertilization increases the frequency of biallelic mutation rate as compared to microinjection^24^. All the lambs generated from CRISPR-Cas9 electroporated embryos contained insertions or deletions at the targeted sites. This 100% mutation rate suggests the efficiency of electroporation as an effective zygote delivery method for CRISPR-Cas9 gene editing and has great potential for the generation of genome edited sheep. Additionally, using this approach, all the edited alleles present in the genome-edited lambs were predicted to result in a KO of the target *SOCS2* gene (Table 1). However, the intended deletion of 85 bp did not occur in any of the founder animals and there was a high degree of variability in the exact mutations that resulted following editing.

Of particular importance to livestock genome engineers and regulators is that 25% of all alleles characterized by WGS were undetectable by PCR and Sanger sequencing of a standard-length amplicon using PCR primers close to the target site. This is consistent with the results of a recent comprehensive analysis of CRISPR/Cas9 genome editing outcomes in human hematopoietic stem and progenitor cells using long-read Pacific Biosciences (PacBio) single-molecule real-time sequencing technology (SMRT-seq) that found large deletions (>200bp) occurred at a frequency of 11.7-35.4% depending on gene target^35^. Similarly, work undertaken in cattle using a dual guide approach also resulted in large deletions ^36^. The mechanism of long deletion is not well understood, but microhomologies are commonly associated with the repair of large deletions and were observed in our results^37^. Large deletions pose a major problem for the genotyping of genome edited livestock embryos where small quantities of DNA limit the possibilities for whole genome sequencing.

One major concern regarding CRISPR/Cas9 genome editing is the potential for off-target mutations. Off-target mutations could hypothetically result in unintended deleterious effects in sheep. The location of sites with 3 or 4 bp mismatches can be predicted effectively using web-based tools like CasOFFinder that search reference genomes for potential off-target sites ^4, 23, 38^. Our analysis of potential off-target sites with 4 mismatches or fewer from the target sequence identified a small number of mutations which may be a result of off-target activity, but notably in-range mutations were also observed in unedited controls (Supplementary Table 4). Further, we observed several orders of magnitude more mutations outside of this region in both edited animals and controls (Figure 5). This highlights that natural mutations occur frequently in unedited food-producing animals (i.e. controls). In this case, the unedited rams had on average of 5.1 million naturally-occurring SNP and indel mutations relative to the sheep reference genome. Similarly, a project that sequenced 2,703 cattle of different breeds found 84 million SNPs and 2.5 million naturally-occurring indels ^39^. Despite this known variation, animals produced by conventional breeding methods are not routinely evaluated for unintended mutations at the molecular level as genetic variation per se is not to be known to be associated with food safety hazards ^40^. Increasing the search for off-target sites from those with up to a 3 bp mismatch (n=4) with the sequence of the two sgRNA guides, to those with up to a 4 bp mismatch (n=133) resulted in a more than 30-fold increase in the number of potential off-target sites to be analyzed. The purpose or food safety value of requiring off-target analyses be performed on the genome of healthy gene-edited sheep is unclear.

Several specialized programs for detection of SV have been developed, such as Manta ^41^, DELLY ^42^, Vg-Graph ^43^, and Minigraph-Cactus ^44^, along with commercial software such as CLC Genomics Workbench (QIAGEN, https://digitalinsights.qiagen.com) among others. However, the only accurate way to identify SV is for species with defined haplotypes, such as with human populations ^44^. This conclusion is based on the assumption that the SV only occurs on one chromosome. Unfortunately, if multiple SV occur in the same region, as with CRISPR edits, preexisting haplotypes are irrelevant for that region. Even worse, if the SVs are at intermediate frequency, i.e. a heterozygote, a consensus sequence for that region is not possible. A work-around was developed for our research whereby the haplotypes were resolved by manual inspection of the read overhangs, extraction of those reads, Denovo realignment, extraction of the consensus sequence, followed by a BLAST to the reference to reconstruct the SV. While this method is highly accurate, the procedure is laborious but tractable for examination of on-site edits. However, the method is not tractable for assessment of off-site edits of 4 or more misalignments. Nevertheless, programs such as CLC Genomics, and perhaps others are accurate at finding where breakpoints have occurred, even if the programs are not accurate in reassembling the results into actual SV. It may be that for the purposes of finding off-target effects, finding breakpoints is adequate for the purpose. However, as discussed previously the value of these analyses is questionable given that genetic variation per se is not to be known to be associated with food safety hazards. The results from this paper support the mounting evidence that off-target editing is rare with well-designed gRNAs.

Another concern in genome editing is mosaicism, which can obscure the evaluation of founder animal phenotypes and result in a variety of offspring phenotypes ^4^. Mosaicism that includes some remaining wildtype sequence is repeatedly observed with engineered nucleases and can diminish the value of single-step genome editing to produce gene edited founders ^45^. This could explain the presence of a low level of *SOCS2* mRNA transcripts and proteins.

The differences in growth between *SOCS2*(-/-) KO and wild type sheep align well with the growth phenotype seen in the *Socs2(-/-)* mice, and they are larger than what has previously been reported in sheep with a naturally-occurring *SOCS2* mutation ^22^. However, while the results are promising, there are still many caveats to these results including the limited sample size, the known effects of sex, litter size and birthweight on growth, and the unknown maternal genetic background of the gene edited sheep given the slaughter-house ovary origin of the oocytes that were fertilized.

The high growth phenotype could potentially be a very valuable trait for introduction into sheep breeding programs, especially through the use of *SOCS2*(-/-) rams as terminal sires. The greatly improved growth phenotype, without an associated increase in birth weight, could decrease the time to market weight, the amount of feed consumed, and reduce the carbon intensity of producing a unit of lamb. However, negative effects observed on reproduction have been observed in *Socs2*(-/-) mice including a failure to maintain pregnancy in certain lines ^46^, and a reduced lifespan ^47^. More data is required to determine how sheep carrying heterozygous mutations in the *SOCS2* gene perform under commercial conditions, and if there are any negative phenotypic correlations with other traits or pleiotropic effects.

## Conclusions

This study documents a comprehensive genotypic analysis of CRISPR/Cas9 genome editing of six genome edited sheep. Our findings demonstrate that electroporation with dual sgRNAs was highly efficient in generating *SOCS2* knockout sheep, with a 100% mutation rate and the ability to processes multiple embryos simultaneously. Notably, exhaustive analysis of potential-off target edits with up to 4 mismatches to the guide RNA did not reveal concerning off-target activity with this approach, which will be highly applicable to other livestock editing endeavors. The growth phenotype of the *SOCS2* (-/-) sheep suggests there are likely benefits for productivity which will be investigated further in the next generation.

## Acknowledgments

We would like to express our deep gratitude to Superior Farms (Dixon, CA) for providing sheep ovaries.

## Author Contributions

A.M, P.R, AVE, and J.M. designed the experiments. A.M. and D.F. carried out the experiment. B.M. and T.U. performed the surgical procedure of embryo transfer. D.H. performed the whole genomic sequencing analysis to visualize on-target edits and assist with primer design. W. M. and N.W. performed both on target confirmation and off-target analysis using WGS data. A.M., D.F. AVE and P.R. wrote the manuscript. N.W. prepared figures, and all authors reviewed the final manuscript.

## Data Availability

The data presented in the study are deposited in the NCBI Sequence Read Archive (SRA) repository, accession numbers SAMN42100813–SAMN42100820. BioProject: PRJNA1128721 (https://www.ncbi.nlm.nih.gov/bioproject/PRJNA1128721) .

## Supplementary Information

**Supplementary Table 1.**
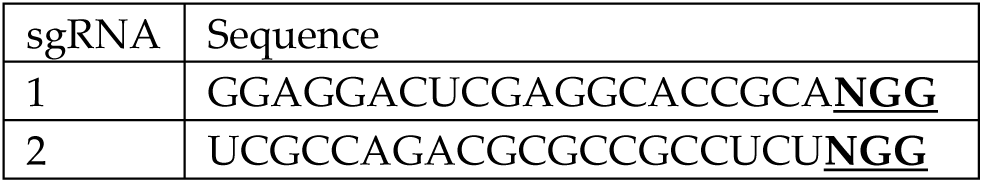
sgRNAs sequences used to target exon 1 of *SOCS2* gene in sheep. PAM site is bold, underlined.

**Supplementary Table 2:**
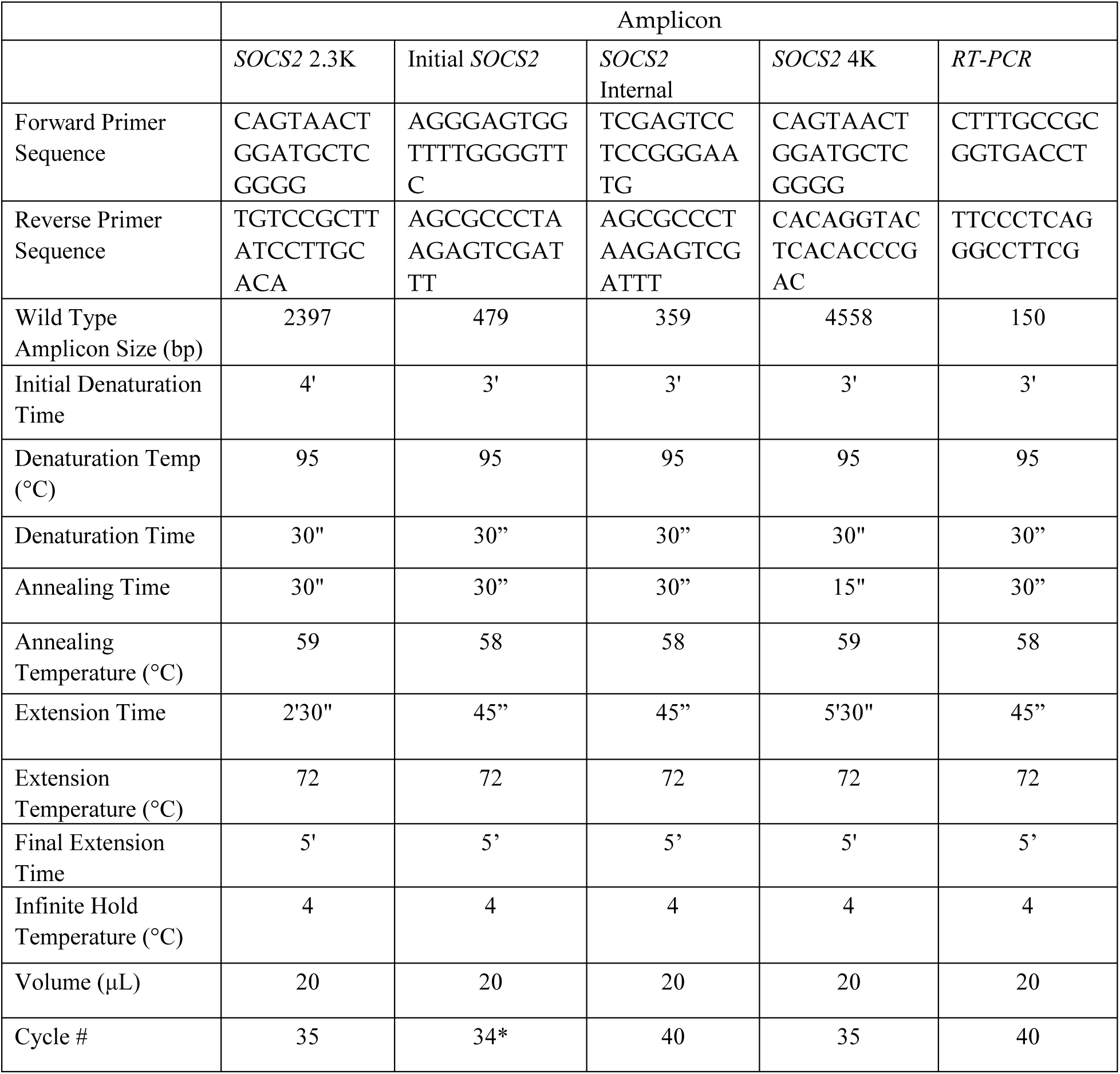
Primer sequences and amplification conditions for PCR of each SOCS2 amplicon.

**Supplementary Table 3.**
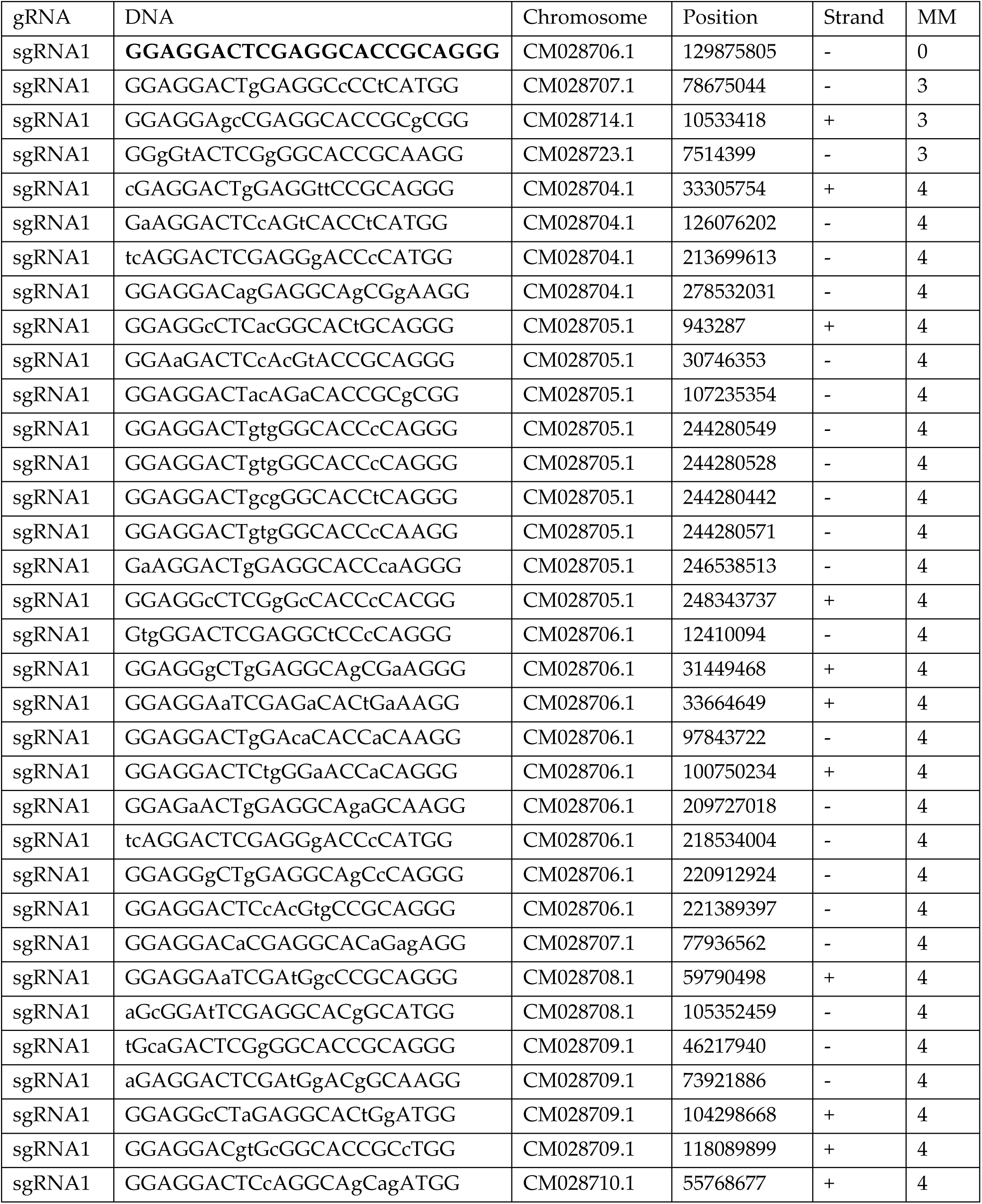
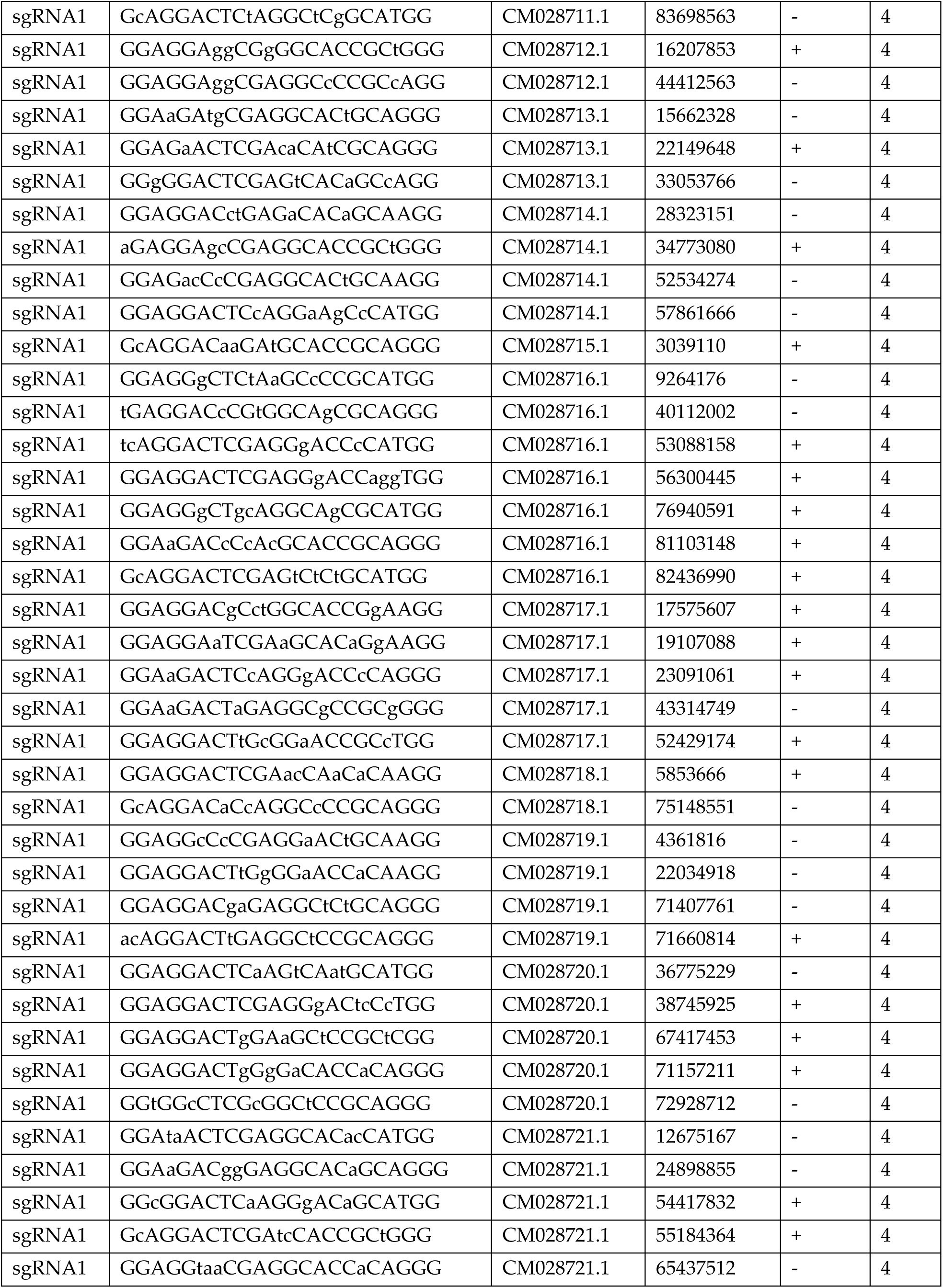
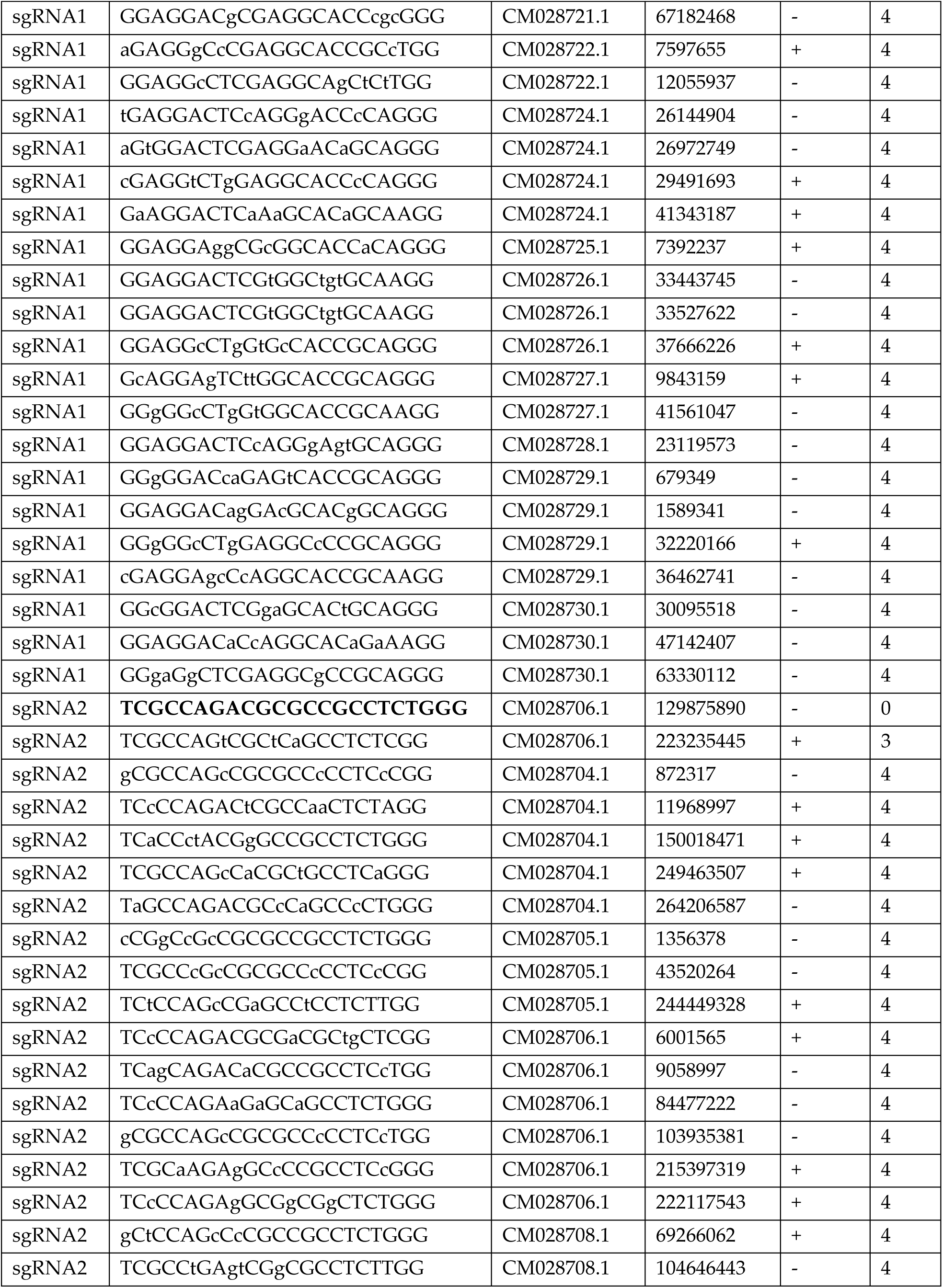
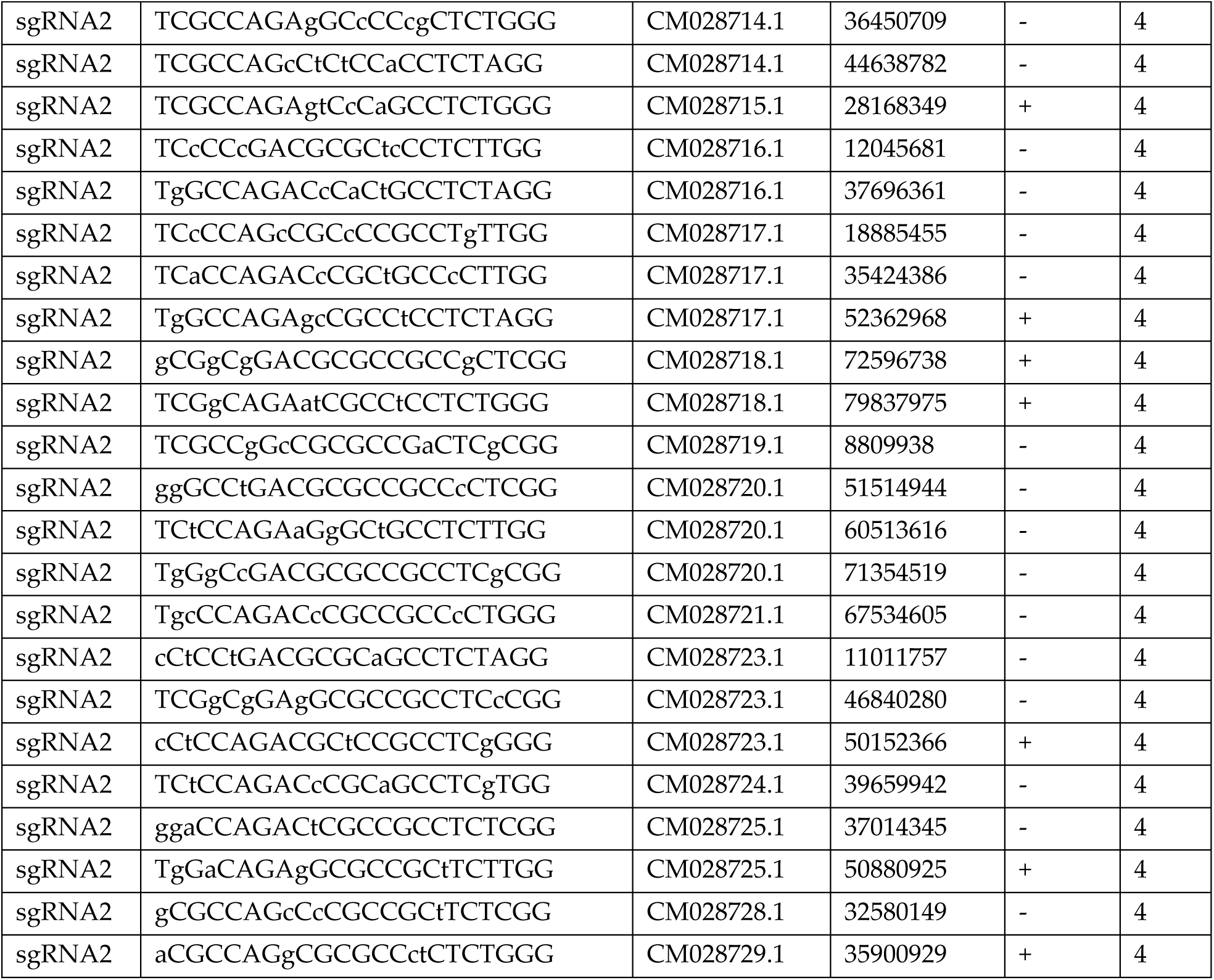
List of potential off-targets for the ARS-UI_Ramb_V2.0 genome. "MM" indicates the number of mismatches between the crRNA and potential target DNA.

**Supplementary Table 4.** Potential off-target mutation identification in six gene edited (7181-7186) and two control (Ram 1-2) sheep. Indel and structural variant mutation data filtered to detect mutations occurring at a region with a 3-4 nucleotides sgRNA mismatched binding site as detailed in Supplementary Table 3 (±100 bp).

| Strand | MM | Chromosome | Mutation Region Start | Mutation Region End | Start Minus 100 bp | End Plus 100 bp | Type | Evidence | Binding Site Within Mutated Region | Binding Site Within 100 bp Of Mutation | Sample |
| --- | --- | --- | --- | --- | --- | --- | --- | --- | --- | --- | --- |
| - | 4 | 18 | 65437258 | 65440611 | 65437158 | 65440711 | Deletion | Cross mapped breakpoints | TRUE | TRUE | 7182 |
| - | 4 | 1 | 126076202 | 126076252 | 126076102 | 126076352 | Insertion | Close breakpoints | TRUE | TRUE | 7184 |
| - | 4 | 26 | 1589265 | 1589665 | 1589165 | 1589765 | Complex | Cross mapped breakpoints (invalid orientation) | TRUE | TRUE | 7185 |
| + | 4 | 14 | 17575557 | 17575607 | 17575457 | 17575707 | Insertion | Close breakpoints | TRUE | TRUE | 7185 |
| - | 4 | 3 | 97843675 | 97843750 | 97843575 | 97843850 | Insertion | Close breakpoints | TRUE | TRUE | 7185 |
| - | 4 | 2 | 244280490 | 244280575 | 244280390 | 244280675 | Deletion | Self mapped | TRUE | TRUE | 7185 |
| - | 4 | 2 | 1356435 | 1356436 | 1356335 | 1356536 | Insertion | Self mapped | FALSE | TRUE | 7181 |
| + | 4 | 5 | 69266102 | 69266103 | 69266002 | 69266203 | Insertion | Self mapped | FALSE | TRUE | 7181 |
| + | 4 | 5 | 69266135 | 69266136 | 69266035 | 69266236 | Insertion | Self mapped | FALSE | TRUE | 7181 |
| - | 4 | 5 | 105352373 | 105352399 | 105352273 | 105352499 | Insertion | Close breakpoints | FALSE | TRUE | 7181 |
| + | 3 | 11 | 10533303 | 10533390 | 10533203 | 10533490 | Insertion | Close breakpoints | FALSE | TRUE | 7183 |
| + | 4 | 10 | 22149689 | 22149690 | 22149589 | 22149790 | Deletion | Self mapped | FALSE | TRUE | 7183 |
| + | 4 | 3 | 215397280 | 215397281 | 215397180 | 215397381 | Insertion | Self mapped | FALSE | TRUE | 7183 |
| - | 4 | 3 | 218533991 | 218534000 | 218533891 | 218534100 | Insertion | Close breakpoints | FALSE | TRUE | 7183 |
| - | 4 | 2 | 1356398 | 1356399 | 1356298 | 1356499 | Insertion | Self mapped | FALSE | TRUE | 7184 |
| - | 4 | 2 | 1356435 | 1356436 | 1356335 | 1356536 | Insertion | Self mapped | FALSE | TRUE | 7184 |
| + | 4 | 23 | 37666258 | 37666267 | 37666158 | 37666367 | Insertion | Close breakpoints | FALSE | TRUE | 7184 |
| - | 4 | 13 | 40111996 | 40111998 | 40111896 | 40112098 | Insertion | Close breakpoints | FALSE | TRUE | 7184 |
| + | 4 | 13 | 56300417 | 56300418 | 56300317 | 56300518 | Deletion | Self mapped | FALSE | TRUE | 7184 |
| - | 4 | 3 | 218533991 | 218534000 | 218533891 | 218534100 | Insertion | Close breakpoints | FALSE | TRUE | 7184 |
| + | 4 | 2 | 943251 | 943278 | 943151 | 943378 | Insertion | Close breakpoints | FALSE | TRUE | 7185 |
| + | 3 | 11 | 10533455 | 10533456 | 10533355 | 10533556 | Insertion | Tandem duplication | FALSE | TRUE | 7185 |
| - | 4 | 13 | 12045596 | 12045640 | 12045496 | 12045740 | Insertion | Close breakpoints | FALSE | TRUE | 7185 |
| - | 4 | 21 | 26144866 | 26144872 | 26144766 | 26144972 | Insertion | Close breakpoints | FALSE | TRUE | 7185 |
| + | 4 | 1 | 33305658 | 33305677 | 33305558 | 33305777 | Insertion | Close breakpoints | FALSE | TRUE | 7185 |
| - | 4 | 2 | 244280490 | 244280575 | 244280390 | 244280675 | Deletion | Self mapped | FALSE | TRUE | 7185 |
| - | 4 | 2 | 1356435 | 1356436 | 1356335 | 1356536 | Insertion | Self mapped | FALSE | TRUE | 7186 |
| + | 4 | 22 | 7392328 | 7392329 | 7392228 | 7392429 | Insertion | Self mapped | FALSE | TRUE | 7186 |
| + | 3 | 11 | 10533285 | 10533390 | 10533185 | 10533490 | Insertion | Close breakpoints | FALSE | TRUE | 7186 |
| - | 4 | 2 | 1356398 | 1356399 | 1356298 | 1356499 | Insertion | Self mapped | FALSE | TRUE | Ram 1 |
| - | 4 | 2 | 1356435 | 1356436 | 1356335 | 1356536 | Insertion | Self mapped | FALSE | TRUE | Ram 1 |
| + | 4 | 14 | 19106969 | 19107028 | 19106869 | 19107128 | Insertion | Close breakpoints | FALSE | TRUE | Ram 1 |
| - | 4 | 5 | 105352398 | 105352399 | 105352298 | 105352499 | Insertion | Tandem duplication | FALSE | TRUE | Ram 1 |
| - | 4 | 2 | 107235421 | 107235441 | 107235321 | 107235541 | Insertion | Close breakpoints | FALSE | TRUE | Ram 1 |
| - | 4 | 3 | 218533992 | 218533999 | 218533892 | 218534099 | Replacement | Paired breakpoint | FALSE | TRUE | Ram 1 |
| - | 4 | 21 | 26972625 | 26972683 | 26972525 | 26972783 | Insertion | Close breakpoints | FALSE | TRUE | Ram 2 |
| + | 4 | 13 | 56300417 | 56300417 | 56300317 | 56300517 | Deletion | Self mapped | FALSE | TRUE | Ram 2 |
| - | 4 | 15 | 75148453 | 75148461 | 75148353 | 75148561 | Insertion | Close breakpoints | FALSE | TRUE | Ram 2 |
| + | 4 | 3 | 215397268 | 215397269 | 215397168 | 215397369 | Insertion | Self mapped | FALSE | TRUE | Ram 2 |
| - | 4 | 3 | 218533991 | 218534000 | 218533891 | 218534100 | Insertion | Close breakpoints | FALSE | TRUE | Ram 2 |

**Supplementary Table 5.**
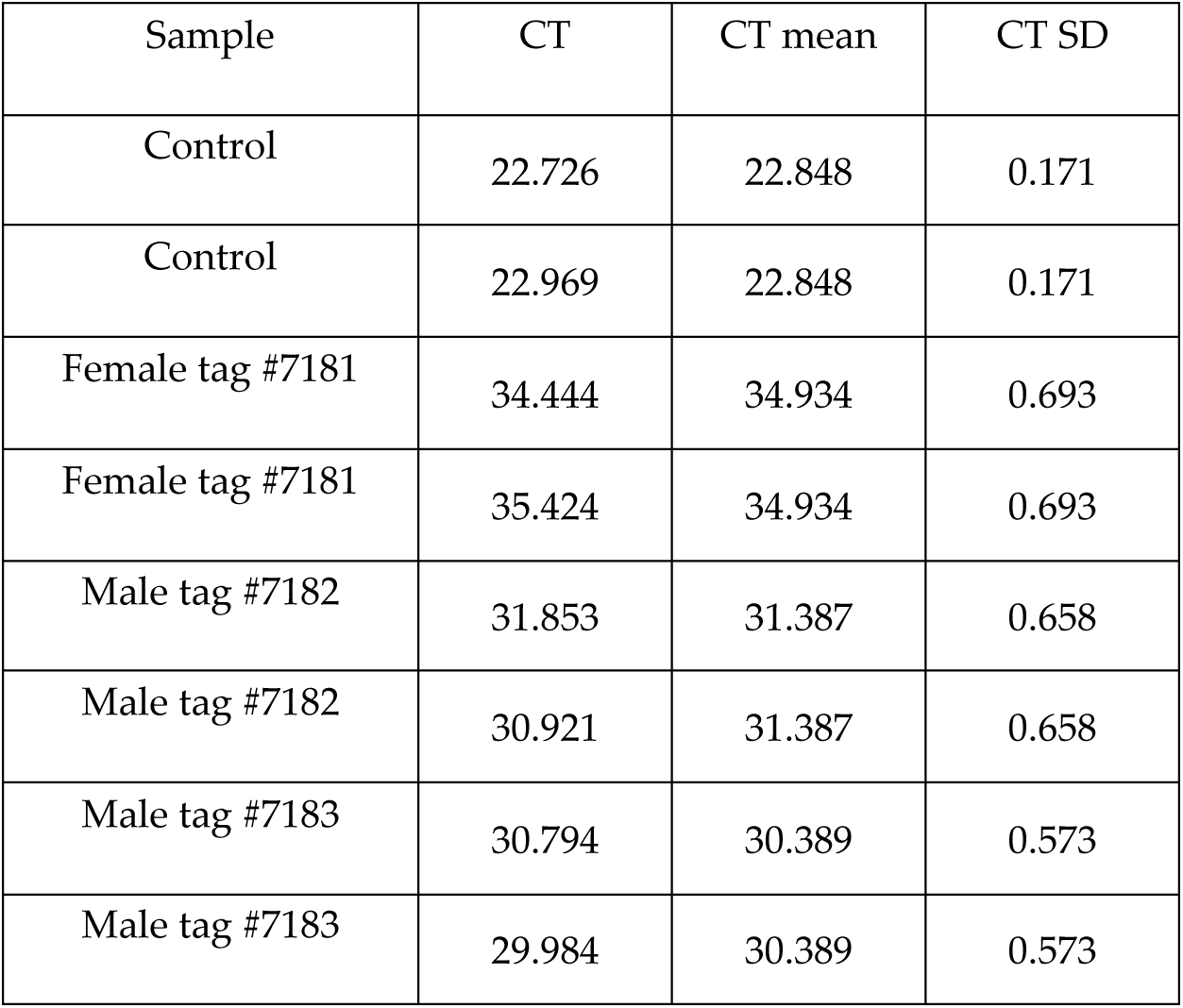
RT-PCR values of mRNA extracted from fibroblast cell lines derive from three healthy *SOCS2* Knock-out lambs produced by zygote electroporation of CRISPR-Cas9.

**Supplementary Figure 1.**
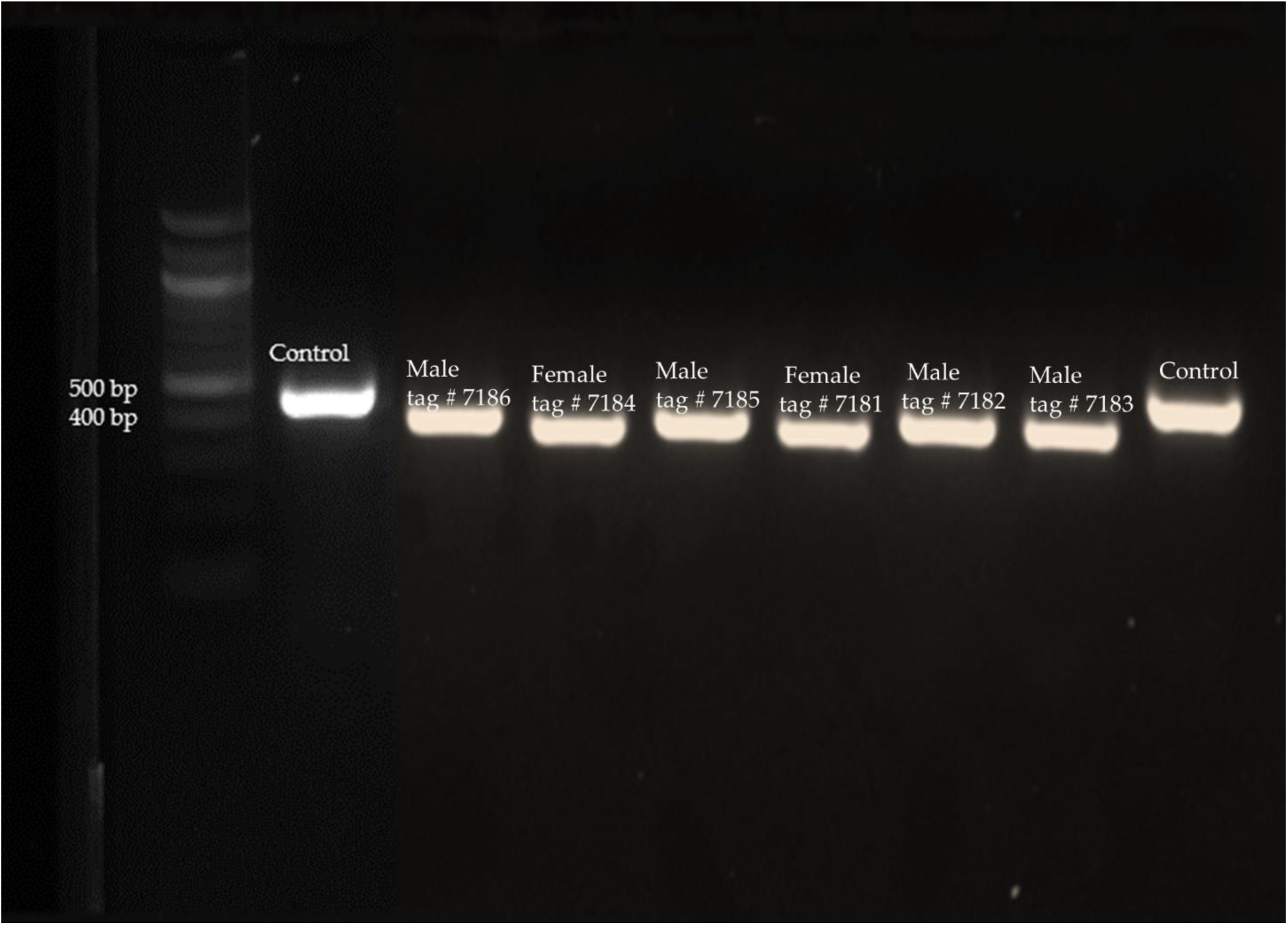
DNA bands of control (479 bp) and SOCS2 knock-out fetuses and lambs when using primers flanking the two target sites in 2% agarose gel.

